# Ki-67 and CDK1 Control the Dynamic Association of Nuclear Lipids with Mitotic Chromosomes

**DOI:** 10.1101/2024.09.10.612355

**Authors:** Hsiao-Tang Hu, Ueh-Ting Tim Wang, Bi-Chang Chen, Yi-Ping Hsueh, Ting-Fang Wang

**Author notes:** Corresponding to Dr. Ting-Fang Wang, and Dr. Yi-Ping Hsueh, Institute of Molecular Biology, Academia Sinica, 128, Sec 2, Academia Rd, Taipei, 11529, Taiwan, ROC.

## Abstract

Nuclear lipids play roles in regulatory processes such as signaling, transcriptional regulation, and DNA repair. In this report, we demonstrate that nuclear lipids may contribute to Ki-67-regulated chromosome integrity during mitosis. In COS-7 cells, nuclear lipids are enriched at the perichromosomal layer and excluded from intrachromosomal regions during early mitosis, but are then detected in intrachromosomal regions during late mitosis, as revealed by TT-ExM, an improved expansion microscopy technique that enables high-sensitivity, super-resolution imaging of proteins, lipids, and nuclear DNA. The nuclear non-histone protein Ki-67 acts as a surfactant to form a repulsive molecular brush around fully condensed sister chromatids in early mitosis, preventing the diffusion or penetration of nuclear lipids into intrachromosomal regions. Ki-67 is phosphorylated during mitosis by cyclin-dependent kinase 1 (CDK1), the best-known master regulator of the cell cycle. Both Ki-67 knockdown and reduced Ki-67 phosphorylation by CDK1 inhibitors allow nuclear lipids to penetrate chromosomal regions. Thus, both Ki-67 protein level and phosphorylation status during mitosis appear to influence the perichromosomal distribution of nuclear lipids. Ki-67 knockdown and CDK1 inhibition also lead to uneven chromosome disjunction between daughter cells, highlighting the critical role of this regulatory mechanism in ensuring accurate chromosome segregation. Given that Ki-67 has been proposed to promote chromosome individualization during open mitosis in vertebrates, our results reveal that nuclear lipid enrichment at the perichromosomal layer enhances Ki-67’s ability to form a protective chromosomal envelope, which is critical for correct chromosome segregation and maintenance of genome integrity during mitosis.

## Introduction

Lipids are key biomolecules that form the structures of biomembranes and serve as signaling molecules to regulate cellular responses. In the nucleus, neutral and polar lipids are found in the nuclear envelope (NE), nucleolus, and nuclear matrix (Ledeen and Wu, 2008, Garcia-Gil and Albi, 2017, Moriel-Carretero, 2021). These lipids can be synthesized *de novo* in the inner nuclear membrane and nuclear matrix or generated at the endoplasmic reticulum connected to the NE and transported into the nucleus (Payrastre et al., 1992, Vann et al., 1997, Romanauska and Köhler, 2018, Fujimoto, 2022). Some nuclear lipids form droplets that serve as nucleolar scaffolding structures, splicing speckles, and DNA repair foci (Osborne et al., 2001, Mortier et al., 2005, Wang et al., 2017). In addition to serving as structural components, nuclear lipids also participate in a variety of biological functions, including in transcriptional regulation (Zou et al., 2011, Yildirim et al., 2013, McDonnell et al., 2016), signaling (Wang et al., 2017, Zhang et al., 2007, Yildirim et al., 2013), the DNA damage response (Wang et al., 2017, Zhang et al., 2007), and protein storage (Cermelli et al., 2006, Kumanski et al., 2021). Therefore, nuclear lipids have multiple activities.

Ki-67, a giant nuclear non-histone protein comprising 3256 amino acids, is a well-known marker of cell proliferation (Andrés-Sánchez et al., 2022). The subnuclear distribution of Ki-67 changes significantly during the cell cycle (Sun and Kaufman, 2018). Ki-67 is concentrated in the nucleolus and in some punctate structures during interphase, but it relocates to transparent gel-like structures surrounding condensed chromosomes (termed the perichromosomal layer or chromosomal periphery) during mitosis (Kill, 1996, Booth and Earnshaw, 2017, Sun and Kaufman, 2018, Gautier et al., 1992). Based on analyses of Ki-67, the perichromosomal layer is thought to isolate (or individualize) sister chromatid pairs from other pairs and to promote efficient interactions with the mitotic spindle during mitosis (Cuylen et al., 2016). Ki-67 also excludes the cytoplasm from condensed chromosomes during mitosis (Cuylen-Haering et al., 2020).

The perichromosomal layer mostly originates from the nucleolus (Booth and Earnshaw, 2017, Ma et al., 2007). When a cell enters mitosis, the nucleolus dissipates and its components relocate to cover the surface of mitotic chromosomes (Booth and Earnshaw, 2017, Ma et al., 2007). In addition to Ki-67, other nucleolar components, including nucleolin and ribosomal RNA, are present in the perichromosomal layer (Hsu et al., 1965, Moyne and Garrido, 1976, Ma et al., 2022). These proteins and RNA molecules form a liquid-like coat on mitotic chromosomes and regulate chromosome behavior during mitosis (Hernandez-Armendariz et al., 2024). Although the nucleolus also contains lipids (Wang et al., 2017, Guillen-Chable et al., 2021, Morovicz et al., 2022), it is currently unclear if nucleolar lipids also relocate to the perichromosomal layer and functionally contribute to this interesting biological process. Thus, whether nuclear lipids are present or even enriched at the perichromosomal layer remains elusive.

Expansion microscopy (ExM) is a super-resolution imaging technology that physically expands biological samples by a factor of four or greater, thereby improving the spatial resolution of optical microscopy (Chen et al., 2015, Wassie et al., 2019, Wang et al., 2023, Hu et al., 2023). The biological sample is first embedded in a hydrogel and then physically crosslinked within it before undergoing protease digestion. The biological sample then expands isotropically with the hydrogel when placed in water (Chen et al., 2015, Wassie et al., 2019). Since its original description, many variations of the ExM protocol have been developed to achieve a resolution comparable to those of electron microscopy, including iterative expansion microscopy (iExM) (Chang et al., 2017), pan-ExM (M’Saad and Bewersdorf, 2020), and iterative ultrastructure expansion microscopy (iU-ExM) (Louvel et al., 2023). However, concerns have arisen regarding the use of these iterative methods to visualize membranes or lipid-containing subcellular structures due to their use of strong lytic and denaturing reagents such as sodium dodecyl sulfate (SDS) and/or sodium hydroxide (NaOH) (Chang et al., 2017, M’Saad and Bewersdorf, 2020, Louvel et al., 2023).

To visualize both proteins and lipids of cultured mammalian cells at a super-resolution scale, we established a modified protocol called TT-ExM that involves application of trypsin and tyramide signal amplification (<u>T</u>SA) technologies. We have demonstrated previously that trypsinization retains convincing fluorescent signals in ExM (Hu et al., 2023, Wang et al., 2023). A combination of a biotin-conjugated lipid, i.e., biotin-DHPE (N-(Biotinoyl)-1,2-dihexadecanoyl-sn-glycero-3-phosphoethanolamine), and TSA technology clearly labeled the membranes and lipid-containing structures of cells, including mitochondria, the nuclear envelope (NE), endoplasmic reticulum, Golgi apparatus, lipid-containing vesicles in the cytoplasm, and lipid-containing foci in the nucleolus (Wang et al., 2023). Notably, we have shown that our system reveals the localization of intranuclear lipid components at the perichromosomal layer of mitotic chromosomes (Wang et al., 2023). Accordingly, we speculate that nuclear lipids are involved in forming the perichromosomal layer. In this report, we deploy TT-ExM to illustrate the ultrastructural organization of the perichromosomal layer by revealing the distributions of lipids and Ki-67 during the cell cycle. Our results indicate that Ki-67 regulates the distribution of nuclear lipids at the perichromosomal layer and intrachromosomal region in a cell cycle-dependent manner. Ki-67 knockdown or reducing Ki-67 phosphorylation by means of a CDK1 inhibitor disrupts the distribution of nuclear lipids at the perichromosomal layer. During mitosis, ki-67 and nuclear lipids may act in concert to segregate or individualize the various sister chromatid pairs.

## Results

### Application of TT-ExM to observe lipid-containing structures and chromosomes

We used Biotin-DHPE and tyramide-Alexa fluor-555 in TT-ExM to label and visualize nuclear lipids in COS-7 cells (Wang et al., 2023). After crosslinking to a hydrogel and trypsinization, the sample swells upon immersion in water. PicoGreen, an ultrasensitive green-fluorescent nucleic acid dye for double-stranded DNA, was used to outline nuclear chromosomal DNA. As previously described (Wang et al., 2023), after 4-fold expansion, both cytoplasmic components and nuclear structures of COS-7 cells expanded isotropically, regardless of their cell cycle stages (**Figure 1A** post-expansion). Lipid-containing structures in the cytoplasm, nuclei (e.g., NEs), and nucleoli were clearly labeled by biotin-DHPE (**Figure 1A**), confirming the feasibility of deploying biotin-DHPE in TT-ExM to reveal cellular lipid structures. Notably, biotin-DHPE signals were present in the peripheral regions of mitotic chromosomes (**Figure 1A**, middle magnified inset), suggesting the presence of lipids in the perichromosomal layer. All images above were viewed using a Zeiss LSM700 confocal laser scanning microscope.

**Figure 1.**
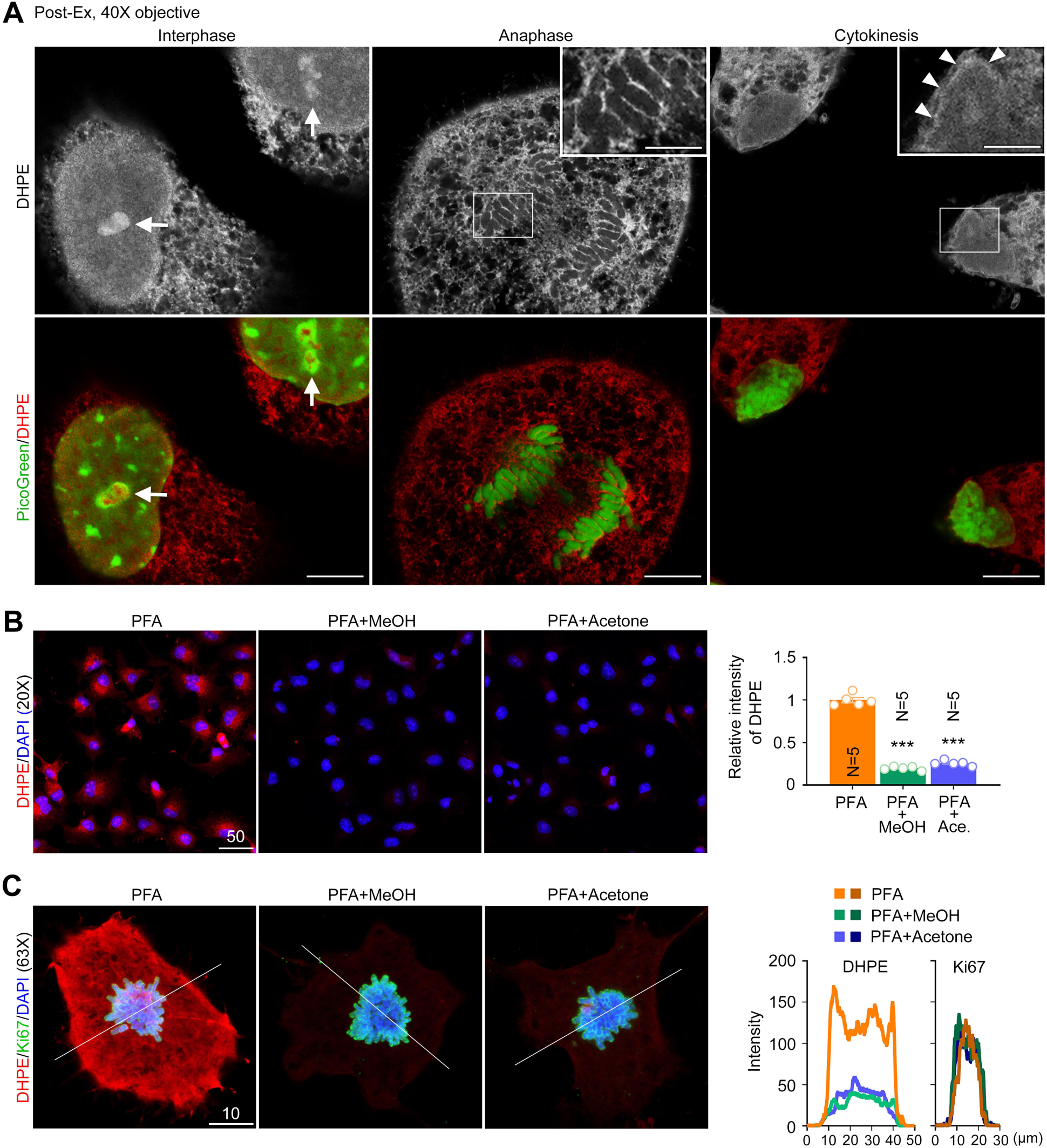
TT-ExM reveals the distribution of lipids and DNA throughout the cell cycle. (**A**) COS-7 cells were stained with biotin-DHPE and PicoGreen and subjected to TT-ExM. Images were acquired using an LSM700 confocal microscope. Different stages of the cell cycle are shown. Arrows indicate the biotin-DHPE signals located in the nucleolus during interphase. The enlarged chromosome segment shows the enriched biotin-DHPE signals at the perichromosomal layers during anaphase. Arrowheads indicate the reforming nuclear envelope after telophase. (**B**) Methanol or acetone treatments reduce biotin-DHPE signals. Left: Representative images. Right: quantification of biotin-DHPE signals. (**C**) Higher magnification images show the effect of methanol or acetone on lipids, but neither treatment affects the signals of Ki-67 or chromosomal DNA. The fine white lines indicate the location of the line scan for quantification shown on the right. Scale bars: (**A**) whole cells: 20 μm; enlarged segment: 8 μm. (**B**) Upper: 50 μm; lower: 10 μm.

To further confirm the lipid signals labeled by DHPE-biotin, we applied organic solvents to extract lipids from paraformaldehyde-fixed cells. After paraformaldehyde fixation and before adding DHPE-biotin, we extracted the lipids by adding methanol or acetone according to methodologies described previously (DiDonato and Brasaemle, 2003, Vore et al., 1974). We found that either the methanol or acetone treatment reduced DHPE-biotin signals by more than 70% when compared to paraformaldehyde fixation alone (**Figure 1B, 1C** upper). In addition to DHPE-biotin staining, DNA counterstaining with 4’,6-diamidino-2-phenylindole (DAPI) and immunofluorescence staining using anti-Ki-67 antibody were also conducted to validate the specific effect of methanol or acetone treatment on lipid extraction (**Figure 1B, 1C**). DAPI is a blue fluorescent DNA stain for which fluorescence is enhanced ∼20-fold upon binding to AT-enriched regions of double-stranded DNA. Ki-67 signals represent nuclear proteins surrounding mitotic chromosomes during cell division. We found that the DHPE-biotin signals were generally reduced in all subcellular regions following the methanol or acetone treatment (**Figure 1B, 1C**). However, the same treatments did not affect chromosomal DNA or nuclear Ki-67 protein (**Figure 1B, 1C**), supporting the specific effect of methanol and acetone in removing lipids. These results confirm that DHPE-biotin indeed specifically labels lipids in cells.

Next, we applied a Zeiss LSM980 confocal laser scanning microscope with an AiryScan 2 area detector to acquire super-resolution images of COS-7 cells stained with DAPI. To quantify the fluorescence signals, we drew a perpendicular line across the chromosome periphery of sister chromatid pairs that had fully condensed during mitosis and performed a line scan analysis of DAPI signals along that line (**Figure 2A**). The expansion index of TT-ExM was determined by comparing the average diameter of fully condensed sister chromatid pairs (**Figure 2B**). Our results show that the average diameters before and after expansion were 1.4 μm and 4.9 μm, respectively, representing an expansion index of 3.5 (**Figure 2C**). Since the LSM980 microscope with AiryScan 2 has a resolution of approximately 90-120 nm, we estimate that the resolution of our TT-ExM protocol is 25-34 nm after 3.5-fold expansion (Hu et al., 2023). Under the same conditions, counterstaining with PicoGreen revealed the highly condensed chromosome axis in the middle of each sister chromatid (indicated by white arrows in **Figure 2D**, right panel). Notably, the surfaces of most, if not all, individual chromatids during mitosis did not appear smooth, suggesting that fully folded chromosomal loops could be observed under our super-resolution conditions (indicated by white asterisks in **Figure 2D**, right panel). These results indicate that TT-ExM is suitable for isotropic expansion of biological macromolecules (e.g., mitotic chromosomes), allowing us to accurately observe the ultrastructures of organelles and subcellular components under an optical microscope.

**Figure 2.**
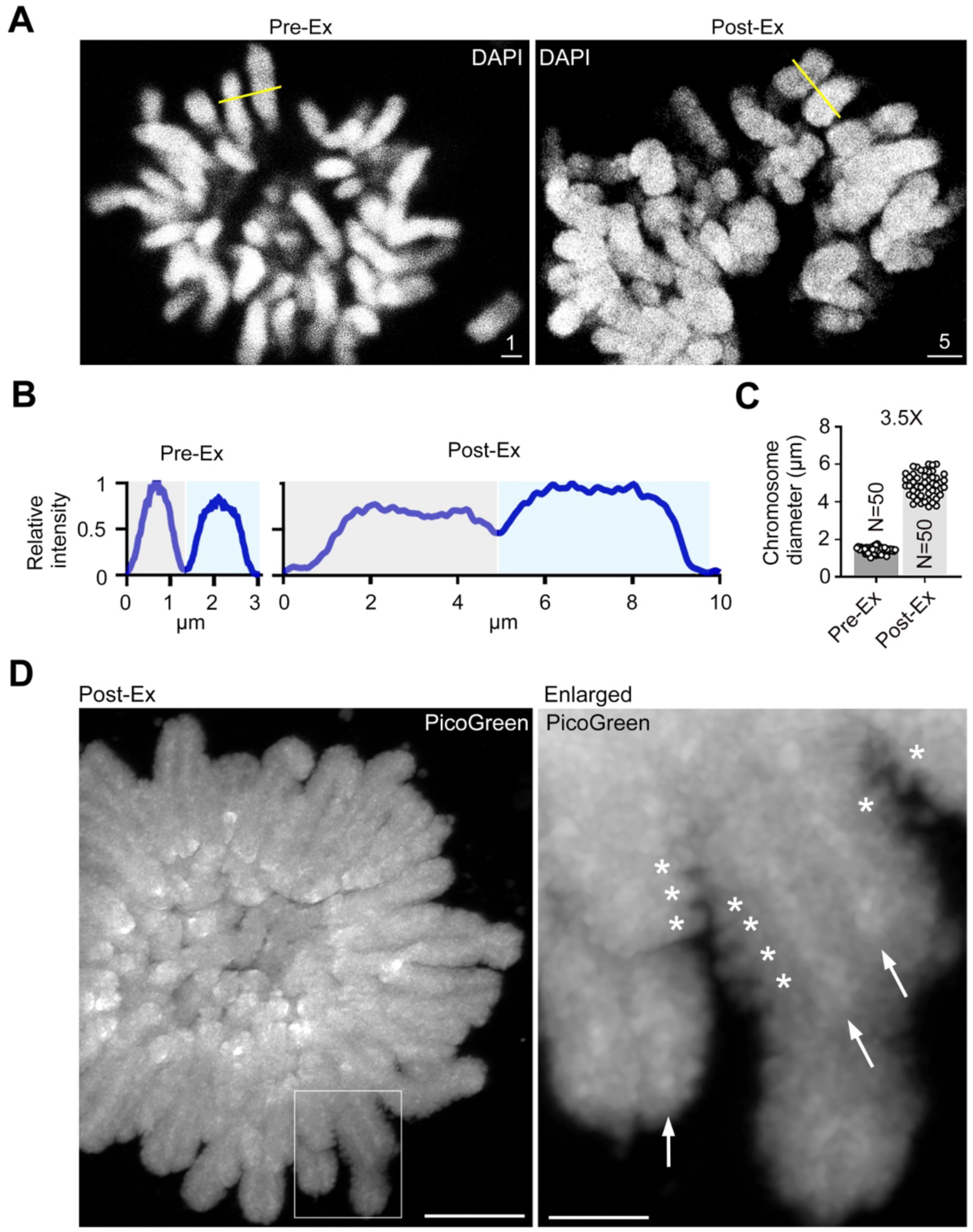
Chromosomal DNA expands isotropically in prometaphase. COS-7 cells were subjected to TT-ExM using PicoGreen or DAPI for DNA staining. Images were acquired using an LSM980 confocal microscope with an Airyscan2 detector. (**A**) Representative images of mitotic chromosomes in prometaphase before and after expansion. Yellow lines are examples of the perpendicular line drawn for line scanning analysis. (**B**)-(**C**) Quantification of chromosome diameters in Pre-Ex and Post-Ex cells. Light grey and blue blocks indicate the regions of two adjacent chromosomes for measurement. Expansion was determined to be 3.5-fold. (**D**) Confocal images of PicoGreen signals in entire Post-Ex cells and enlarged chromosome segments. PicoGreen signal revealed likely chromosome axes displaying a denser DNA signal (arrows), as well as chromosomal loops (asterisks). Scale bars: (**A**) pre-Ex: 1 μm; post-Ex: 5 μm; (**B**) whole cells: 10 μm; enlarged segment: 2.5 μm.

### Dynamic distribution of nuclear lipids during mitosis

We further investigated the distribution of lipid components at the perichromosomal layers. In addition to biotin-DHPE, we dual-stained the cells with an anti-Ki-67 antibody (**Figure 3**) and an anti-phosphatidylserine (PS) antibody (**Figure 4**). Ki-67 is a critical component in forming and organizing the perichromosomal layers (Cuylen et al., 2016, Cuylen-Haering et al., 2020, Ma et al., 2022). It also controls chromosome individualization and genome stability during mitosis (Takagi et al., 2014, Takagi et al., 2018, Garwain et al., 2021, Stamatiou et al., 2024). PS is an anionic glycerophospholipid. Immunofluorescence signals of anti-PS antibodies specifically localize in epichromatin throughout the cell cycle and are associated with nucleosome core histones (Prudovsky et al., 2012). In fixed and permeabilized cells, epichromatin can be identified by partial co-immunolocalization of Ki-67 with staining signals of particular anti-nucleosome or anti-PS antibodies (Gould et al., 2017).

**Figure 3.**
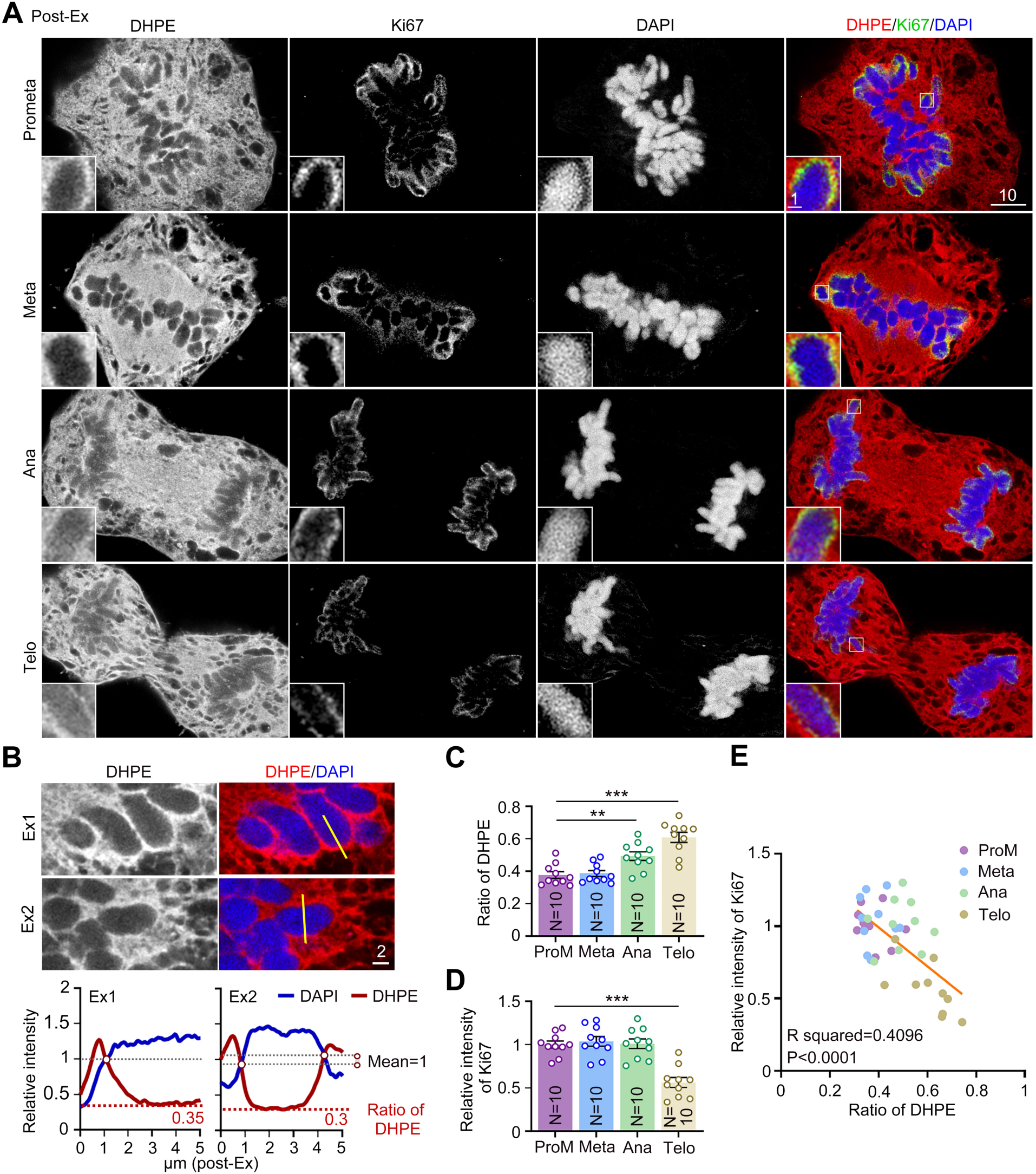
Distribution patterns of Ki-67 and biotin-DHPE signals during mitosis. COS-7 cells were subjected to TT-ExM and stained using biotin-DHPE, Ki-67 antibody and DAPI. Images at different mitotic stages were acquired using an LSM980 confocal microscope with Airyscan2. (**A**) Confocal images of entire cells and enlarged chromosome segments at different stages of mitosis: Prometa (prometaphase); Meta (metaphase); Ana (anaphase); Telo (telophase). The original regions of the enlarged images are indicated only in the rightmost panels. (**B**) Line scanning analysis of biotin-DHPE distribution across the perichromosomal layer and chromosome. Yellow lines indicate the regions analyzed in the two examples, Ex1 and Ex2. Lower panel: The results of line scanning analysis on Ex1 and Ex2. The red open circles indicate the intersection of biotin-DHPE signal and the chromosome boundary, which was set as a ratio of 1. The lowest ratio of biotin-DHPE was identified as the relative lipid level in the chromosome interior. (**C**) Ratio of the biotin-DHPE signal in the intrachromosomal area to that at the periphery. The ratio was lower in prometaphase and metaphase, but increased in anaphase and telophase. (**D**) Relative levels of total Ki-67 immunoreactivity, which clearly declined during telophase. (**E**) Negative correlation of the biotin-DHPE ratio and Ki-67 signal intensity. The sample size “N” indicates the number of examined cells. The data represent mean ± SEM and individual data points are also shown. All data were collected from at least two independent experiments. ** *P* < 0.01; *** *P* < 0.001; one-way ANOVA. Scale bars: (**A**) whole cells: 10 μm; enlarged segment: 1 μm; (**B**) 2 μm.

**Figure 4.**
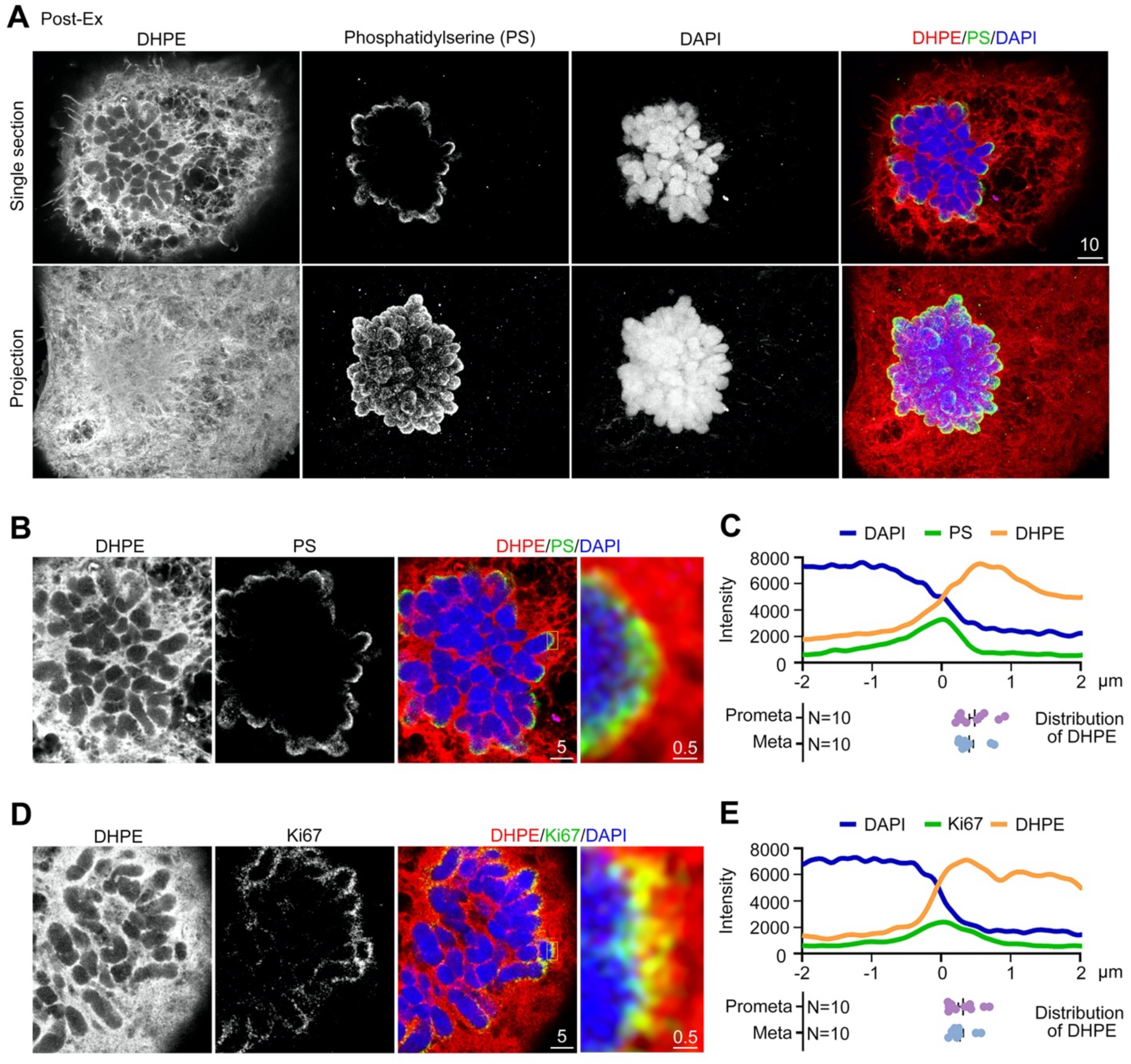
Relative distribution patterns of Ki-67, biotin-DHPE and PS. COS-7 cells were subjected to TT-ExM and stained using antibodies recognizing Ki-67 and phosphatidylserine (PS), as well as for biotin-DHPE and DAPI. Images of cells in prometaphase and metaphase were acquired using an LSM980 microscope with an AiryScan2 detector. (**A**) Confocal images of lipid components (biotin-DHPE and PS) during prometaphase. Upper: Single section. Lower: Z-projection. Biotin-DHPE was distributed throughout cells and covered the entire chromosomes. PS was only located in regions of epichromatin. (**B**)-(**C**) Distribution of biotin-DHPE and PS. (**D**)-(**E**) Distribution of biotin-DHPE and Ki-67. (**B**), (**D**) Representative confocal images. The 10X enlarged images of the insets at regions of epichromatin in the whole-cell images are shown in the rightmost panels. (**C**), (**E**) The results of line scanning. Upper: an example of the line scanning result. Lower: the relative positions of the peaks of biotin-DHPE signals. The peaks of PS and Ki-67 signals at the chromosome edge are set as 0. Biotin-DHPE signals are always present at the outer part of perichromosomal layer relative to the PS and Ki-67 signals. The sample size “N” indicates the number of examined cells in prometaphase and metaphase. Scale bars: (**A**) 10 μm; (**B**), (**D**) whole cells: 5 μm; enlarged segment: 0.5 μm.

We observed that the immunofluorescence signals of biotin-DHPE indeed overlapped with those of Ki-67 at the perichromosomal layers during prometaphase and metaphase (**Figure 3A**). As mitosis progressed into anaphase and telophase, we detected enhanced biotin-DHPE immunofluorescence signals in the chromosomal interior regions. These results indicate that DHPE-labeled nuclear components (e.g., nuclear lipids) had migrated from the perichromosomal layers toward chromosomal cores as mitosis progressed (**Figure 3A**).

To quantify the movement of the biotin-DHPE signals toward the chromosomal interior, we performed line scanning as described in Figure 2 to determine the ratio between the lowest biotin-DHPE signal within chromosomes and the biotin-DHPE signal at chromosome edges, i.e., the inner/edge ratio of DHPE signal (**Figure 3B**). Our results show that this ratio is less than 0.4 during prometaphase and metaphase, but increases to ∼0.5 and ∼0.6 for anaphase and telophase, respectively (**Figure 3C**). By quantifying average global Ki-67 immunofluorescence signals in mitotic cells, we also detected that intracellular Ki-67 levels decline as mitosis progresses from prometaphase to telophase (**Figure 3D**). Given that we uncovered a significant negative correlation between intracellular Ki-67 protein levels and the inner/edge ratio of DHPE signals (**Figure 3E**), we speculate that Ki-67 may prevent DHPE-labeled nuclear components (e.g., nuclear lipids) from moving into chromosomal interiors during prometaphase and metaphase, thereby promoting chromosome individualization.

### Perichromosomal lipid composition and layer organization

Consistent with the previous study (Prudovsky et al., 2012), we also found that PS immunoreactivity specifically localized in epichromatin after specimen expansion (**Figure 4A**, upper). Z-projection images revealed that PS signal specifically occurred on the surface of the mitotic chromosome cluster during prometaphase (**Figure 4A**, lower). This pattern differs from that of biotin-DHPE signals, which surrounded the mitotic chromosomes (**Figure 4A and 4B**). However, biotin-DHPE signals overlapped with the PS signals within the epichromatin region (**Figure 4B**). Our line scan analysis indicated that the peak of PS signals was closer to the edge of mitotic chromosomes compared to the peak of biotin-DHPE signals (**Figure 4C**). Taken together, these results indicate that, in addition to PS, other lipid components are also enriched around chromosomes.

We also analyzed the relative positions of the signal peaks for Ki-67 and biotin-DHPE. Similar to our comparison of PS and biotin-DHPE, we found that the peak of Ki-67 signals was closer to the edge of mitotic chromosomes compared to the peak of biotin-DHPE signals (**Figure 4D, 4E**). Thus, the DHPE-labeled lipid layer appears to be located outside of the Ki-67 layer of the perichromosomal zone. This scenario is consistent with our hypothesis that Ki-67 may prevent inward movement of lipid components into the interior of mitotic chromosomes.

### Ki-67 knockdown causes chromosome aggregation, slows cell cycle progression, and alters the pattern of the microtubule array

To further confirm the role of Ki-67 in controlling lipid distribution across chromosomes and the perichromosomal layer, we studied the effect of Ki-67 knockdown in COS-7 cells. To establish and characterize Ki-67 knockdown, we designed a plasmid expressing an artificial miRNA, Ki67-miR, to downregulate endogenous Ki-67 in COS-7 cells. Immunostaining and immunoblotting revealed that Ki67-miR significantly reduced Ki-67 protein levels in COS-7 cells relative to the control construct Ctrl-miR (**Figure 5A-5C**). Consistent with previous studies (Cuylen et al., 2016, Cuylen-Haering et al., 2020), Ki-67 knockdown prompted mitotic chromosome collapse and aggregation (**Figure 5A**, as revealed by DAPI counterstaining). Quantification analysis revealed that the area occupied by chromosomes was diminished in Ki-67 knockdown cells compared to cells expressing Ctrl-miR (**Figure 5D**). These results demonstrate that we had successfully knocked down Ki-67 and impaired the spatial arrangement of mitotic chromosomes in COS-7 cells.

**Figure 5.**
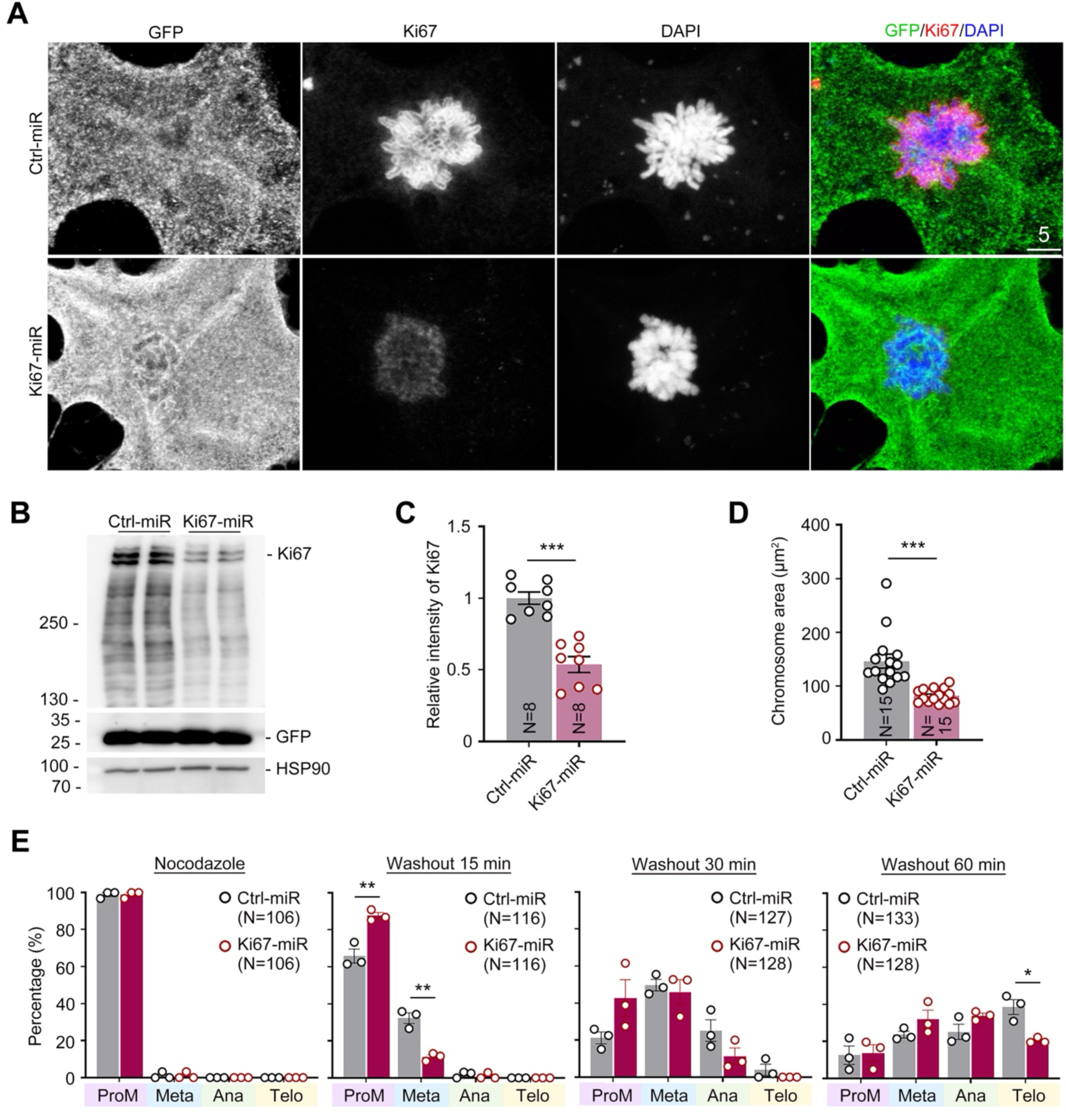
Ki-67 knockdown alters chromosome structure and delays mitotic progress. COS-7 cells were transfected with Ki67-miR or Ctrl-miR and then stained with Ki-67 antibody and DAPI. Since miR constructs coexpress GFP, GFP signals were used to label transfected cells. Images were acquired using an LSM700 confocal microscope. (**A**) Representative confocal images in prometaphase. Ki-67 signal intensity (**C**) and chromosome areas (**D**) were analyzed for transfected cells. (**B**) The knockdown effect of Ki67-miR. Whole-cell extracts were collected and immunoblotted with Ki-67 antibody to detect endogenous Ki-67 protein levels. GFP and HSP90 are loading controls. (**C**) The relative Ki-67 signals in Ki67-miR- and Ctrl-miR-expressing cells. (**D**) Loss of Ki-67 reduces chromosome area. (**E**) Ki-67 knockdown delays progress of mitosis. Cells were treated with 50 nM nocodazole for 6 h to synchronize the cell cycle at prometaphase. After removing nocodazole, the cells were allowed to recover for 15, 30 or 60 mins before being harvested for immunostaining. Mitotic stages were determined according to chromosome patterning. The sample sizes (N) of independent preparations (**C**) or examined cells (**D**, **E**) are indicated in the panels. In (**E**), total cell numbers were collected from three independent experiments. The data represent mean ± SEM and individual data points are also shown. All data were collected from at least two independent experiments. * *P* < 0.05; ** *P* < 0.01; *** *P* < 0.001; two-tailed unpaired Student’s *t*-test. Scale bars: (**A**) 5 μm.

Ki-67 knockdown also delayed the progress of mitosis. First, we applied nocodazole to arrest control or Ki-67 knockdown COS-7 cells at prometaphase (**Figure 5E**, left panel). After washing away nocodazole and allowing the cells to recover for 15, 30, or 60 minutes, we then counted the numbers of COS-7 cells at different stages of mitosis (i.e., prometaphase, metaphase, anaphase, and telophase). Within 15 minutes of washing away nocodazole, more than 30% of the control cells progessed from prometaphase into metaphase. By comparison, only 10% of the Ki-67 knockdown cells entered metaphase under the same treatment (**Figure 5E**). Furthermore, less than 10% of Ctrl-miR- and Ki67-miR-expressing cells remained in prometaphase 60 minutes after nocodazole had been washed away, with ∼40% of the Ctrl-miR-expressing cells entering telophase. In contrast, only 20% of Ki67-miR-expressing cells progressed to telophase. We detected a slight increase in the percentages of Ki-67 knockdown cells in metaphase and anaphase (**Figure 5E**, right). Therefore, Ki-67 knockdown delays mitotic progression.

In addition to chromosomes, we also monitored microtubule patterns in cells expressing Ki67-miR. Since microtubule patterns change during the different stages of mitosis, we focused on the microtubule array in early prometaphase for our comparison (**Figure 6**). At this stage of mitosis, microtubules form a star-like pattern in the center of the chromosome cluster. We observed microtubule filaments extending from the center to the periphery of the chromosome cluster in control COS-7 cells (**Figure 6A**, upper). However, mitotic chromosomes collapsed in Ki67-miR-expressing cells, so although microtubules still formed a star-like pattern in the center of the chromosome cluster of those cells, the star-like pattern was smaller and more irregular than for control counterparts (**Figure 6A**, lower). To quantify the differences, we performed Sholl analysis (Sholl, 1953) to analyze the complexity of microtubules in early prometaphase of control and Ki-67 knockdown cells (**Figure 6B**). Our Sholl analysis revealed that the number of intersections was greatly reduced in Ki-67 knockdown cells (**Figure 6C**), i.e., less than 50% of that enumerated for control cells (**Figure 6D**). We further counted all microtubule filament tips to represent the total number of microtubule filaments in each array, which revealed ∼50% fewer microtubule tips in Ki-67 knockdown cells compared to control cells (**Figure 6E**). Thus, Ki-67 knockdown induces microtubule array abnormalities during mitosis.

**Figure 6.**
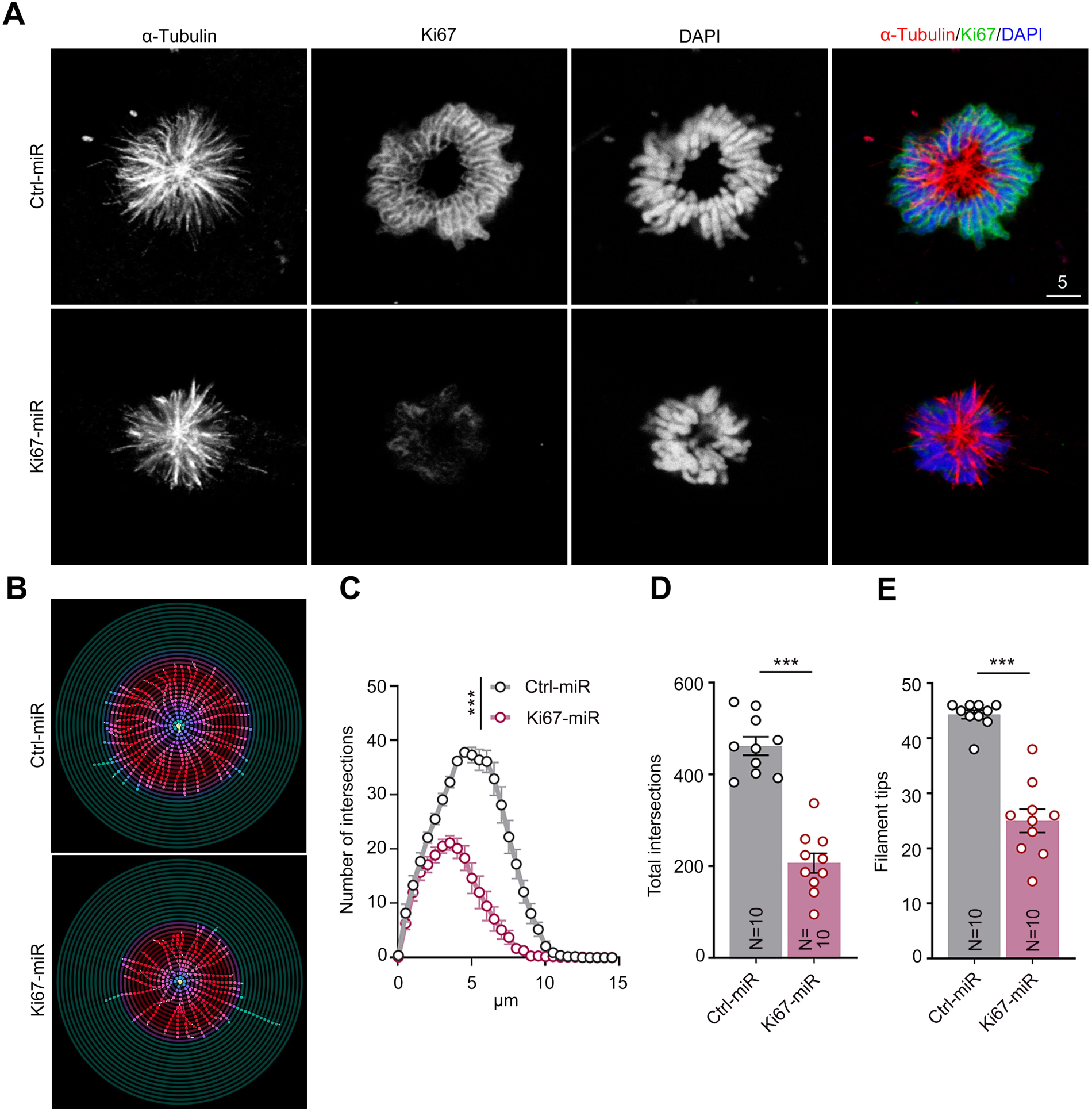
Loss of Ki-67 disturbs microtubule organization. COS-7 cells were transfected with Ki67-miR or Ctrl-miR plasmids and stained with Ki-67 and α-tubulin (for microtubule) antibodies and DAPI. Images were acquired using an LSM700 confocal microscope. (**A**) Confocal images of microtubule filaments during prometaphase. Representative images of Crtl-miR- and Ki67-miR-expressing cells are shown. (**B**)-(**E**) Quantification of microtubule organization by Sholl analysis. (**B**) Representative images of Sholl analysis. (**C**) The distribution of numbers of intersections. (**D**) Total numbers of intersections. (**E**) Total numbers of microtubule filament tips. The sample size “N” indicates the number of examined cells. The data represent mean ± SEM and individual data points are also shown. All data were collected from two independent experiments. *** *P* < 0.001; two-tailed unpaired Student’s *t*-test. Scale bars: **(A)** 5 μm.

Consequently, Ki-67 is critical for chromosome elongation and individualization during mitosis, both of which are required for microtubule attachment to chromosomes to enable mitotic progression.

### Ki-67 knockdown promotes lipid invasion into condensed chromosomes

Lastly, we applied our Ki-67 knockdown system to investigate if Ki-67 is involved in the inward movement of lipids into the chromosome interior during mitosis. Since we detected biotin-DHPE signals as being lowest within condensed chromosomes during prometaphase compared to other mitotic stages (**Figure 3**), we solely analyzed the effects of Ki-67 knockdown during prometaphase. Given that our miR expression construct coexpresses green fluorescent protein (GFP), we used GFP signal to identify transfected cells in culture (**Figure 7A**). Next, we performed line scans to determine biotin-DHPE signals across chromosomes and the perichromosomal layer. GFP signal was included as a control (**Figure 7B, 7C**). We found that the GFP inner/edge ratio was identical (∼0.2) for cells expressing either Ctrl-miR or Ki67-miR (**Figure 7D**), indicating that GFP distribution is insensitive to Ki-67 knockdown. In terms of biotin-DHPE signals, the ratio for control cells was ∼0.28, whereas this ratio increased to 0.34 for Ki-67 knockdown cells (**Figure 7D**), indicating that Ki-67 regulates lipid invasion into condensed chromosomes during mitosis.

**Figure 7.**
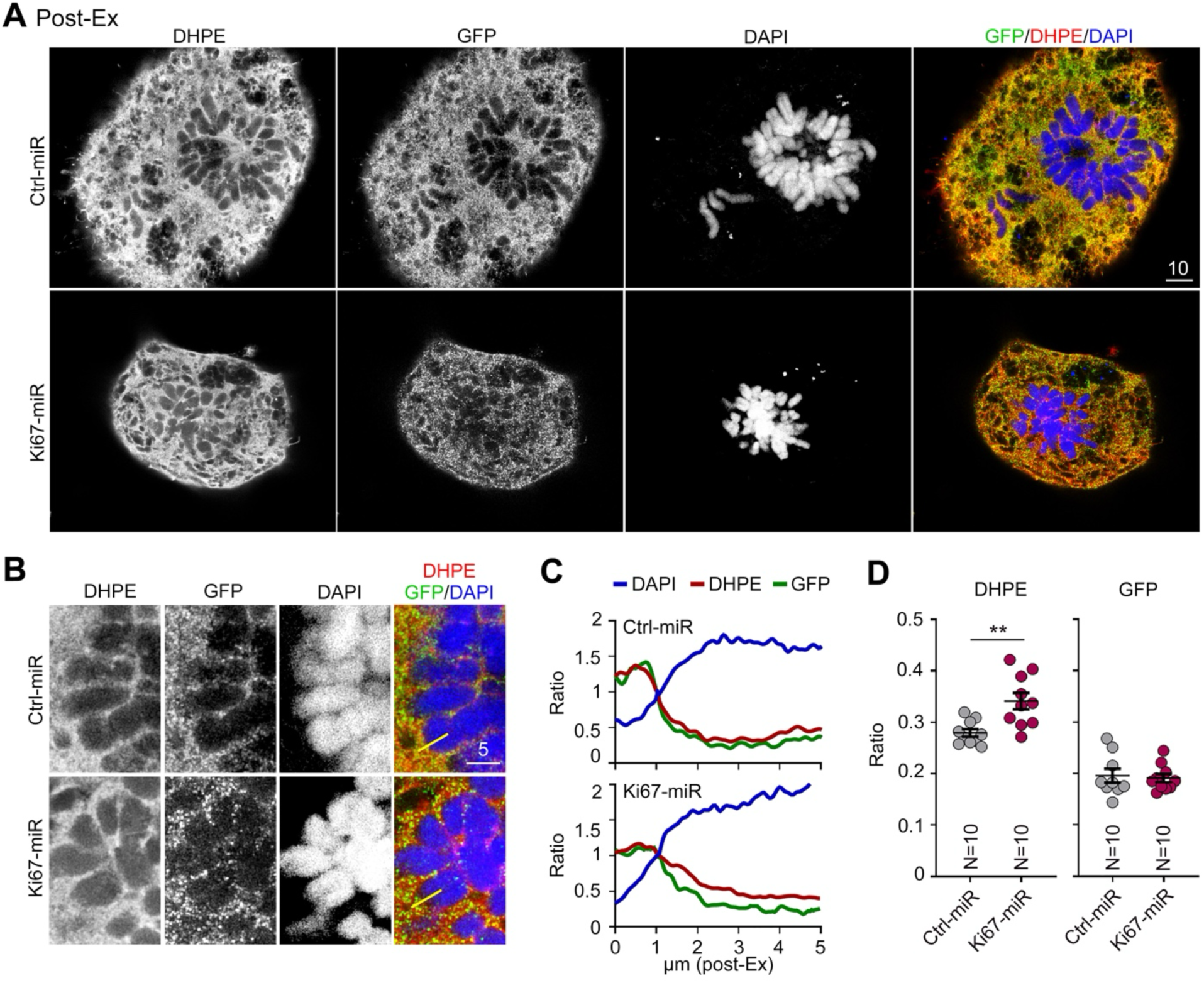
Lipid distribution is altered by Ki-67 knockdown. COS-7 cells were transfected with Ki67-miR or Ctrl-miR plasmids and then stained with biotin-DHPE, Ki-67 antibody and DAPI for TT-ExM. An LSM980 microscope with an AiryScan2 detector was used to acquire images of the cells in prometaphase. (**A**) Representative confocal images. (**B**)-(**D**) Line scanning analysis reveals a change in biotin-DHPE distribution. The yellow lines indicate the regions subjected to line scanning. GFP serves as a control that is insensitive to Ki-67 knockdown. The sample size “N” indicates the number of examined cells. The data represent mean ± SEM and individual data points are also shown. All data were collected from at least two dependent experiments. ** *P* < 0.01; two-tailed unpaired Student’s *t*-test. Scale bars: (**A**) 10 μm; (**B**) 5 μm.

### Blockade of CDK1 activity influences Ki-67 phosphorylation, nuclear lipid distribution and chromosome segregation

To further investigate the relationship among the cell cycle, Ki-67 and nuclear lipid distribution, we applied RO-3306, a specific inhibitor of cyclin-dependent kinase 1 (CDK1), to our culture (**Figure 8A**). CDK1 is an essential protein kinase for cell division (Diril et al., 2012, Massacci et al., 2023). It heavily phosphorylates Ki-67 during mitotic entry (Saiwaki et al., 2005, Takagi et al., 2014, Stamatiou and Vagnarelli, 2021), and regulates Ki-67 localization, especially at the perichromosomal layer, during mitosis (Booth et al., 2014, Booth and Earnshaw, 2017, Remnant et al., 2021, Yamazaki et al., 2022). Our RO-3306 treatment efficiently reduced Ki-67 phosphorylation levels during mitosis (**Figure 8B**). Next, we added RO-3306 into COS-7 cell culture that had been arrested at prometaphase by means of nocodazole (**Figure 8A**). Compared with vehicle control, the RO-3306 treatment altered mitotic progression after nocodazole washout, with the percentages of cells that stayed at prometaphase or anaphase increasing after RO-3306 treatment for 30 min (**Figure 8C**). In contrast, fewer cells were at metaphase upon RO-3306 treatment (**Figure 8C**). These results confirm the effect of CDK1 on Ki-67 phosphorylation and mitotic progression.

**Figure 8.**
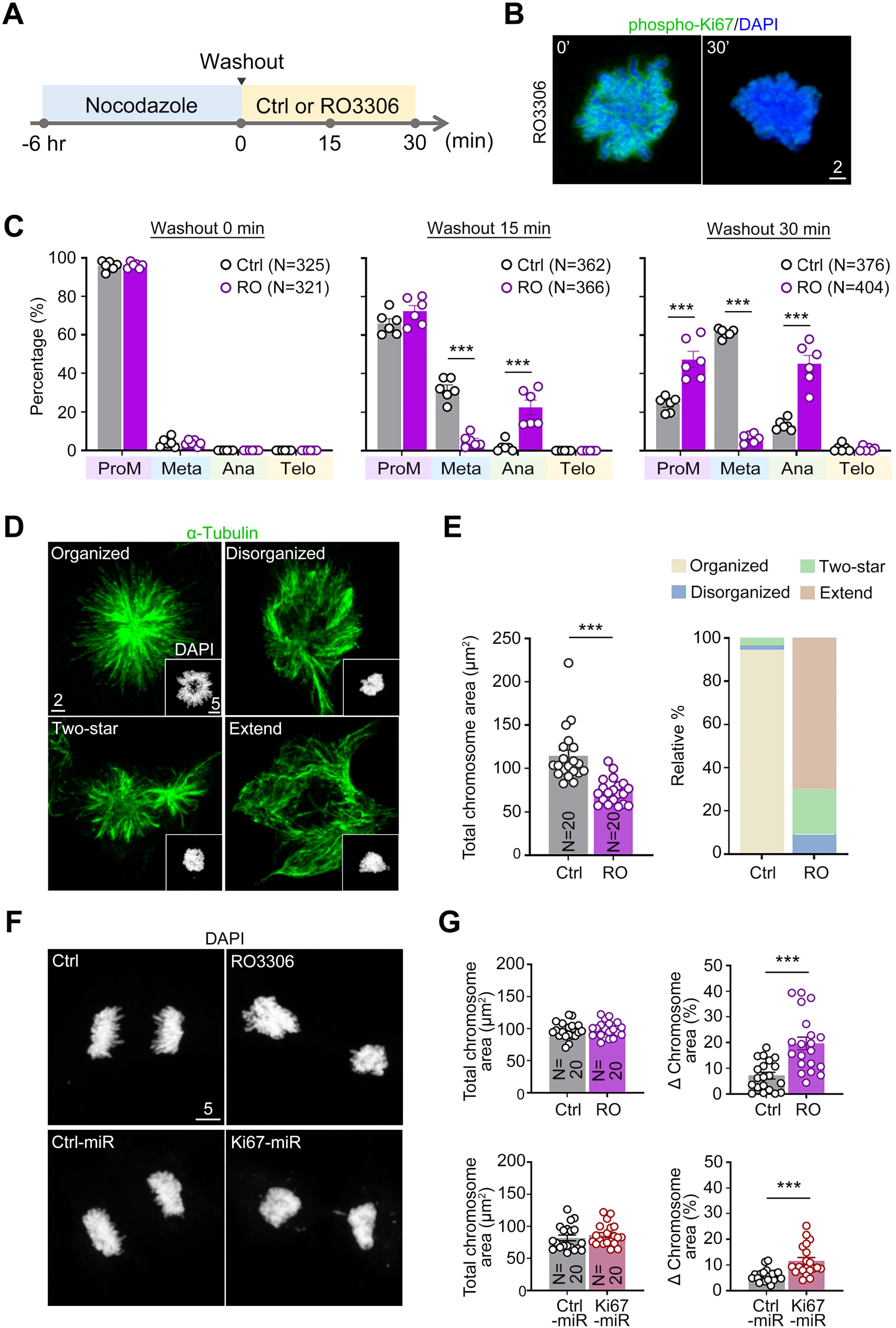
A CDK1 inhibitor reduces Ki-67 phosphorylation, arrests mitotic progress and impairs microtubule organization and sister chromatin separation. (A) Experimental flowchart. Cells were treated with 50 nM nocodazole for 6 hrs to synchronize the cell cycle at prometaphase. After removing nocodazole, cells were treated with 2 μM RO-3306 for 15 or 30 min and then harvested for immunostaining. Vehicle DMSO was included as a control (Ctrl). Particular chromosome patterns determined mitotic stages. (B) RO-3306 treatment for 30 min reduces Ki-67 phosphorylation. (C) RO-3306 treatment disrupts mitotic progress. The percentage of cells in metaphase was reduced, but the proportions of cells in prometaphase and anaphase were increased. (D-E) Reduced Ki-67 phosphorylation disrupts chromosome and microtubule organization. (D) Confocal images of chromosomes and microtubules at prometaphase. Representative images of four types of Ctrl and RO-3306-treated cells are shown. (E) Quantification of chromosome area and microtubule organization patterns. (F-G) Reduced Ki-67 phosphorylation and Ki-67 protein levels both influence sister chromatin separation. (F) Confocal images of chromosome arrangement in anaphase cells. Representative images showing the effects of RO-3306 treatment and Ki-67 knockdown are shown. (G) Quantification of chromosome area and the normalized difference in chromosome area between two daughter cells. The sample size “N” indicates the number of examined cells. The data represent mean ± SEM and individual data points are also shown. All data were collected from two independent experiments. *** *P* < 0.001; unpaired two-tailed *t*-test. Scale bars: (C) whole cells: 2 μm; segment: 5 μm; (E) 5 μm.

Similar to Ki-67 knockdown, we found that chromosomes were collapsed and aggregated after RO-3306 treatment (**Figure 8D**, inset). RO-3306 not only resulted in a reduction in total chromosome area (**Figure 8E**, left), but also altered microtubule organization. In contrast to Ki-67 knockdown that reduced the number and size of microtubule arrays at prometaphase (**Figure 6**), RO-3306 treatment dramatically reduced the number of cells displaying the typical organized star-shaped microtubule arrays, with many of the remaining cells instead displaying extended, two-star, or disorganized microtubule arrays (**Figure 8D, 8E**).

Consistent with our observations of microtubule disorganization, we found that RO-3306 treatment resulted in uneven segregation of chromosomes into the two daughter cells (**Figure 8F, 8G**, upper). This effect of RO-3306 treatment was irrelevant to DNA synthesis since the total chromosome area of both daughter cells was comparable between control and RO-3306-treated cells (**Figure 8G**, upper left). The difference in chromosome area between the two daughter cells was noticeably higher in the RO-3306-treated group compared to the control group (**Figure 8G**, upper right). In Ki-67 knockdown cells, we also observed a similar phenotype in that Ki-67-expressing cells exhibited uneven chromosome segregation (**Figure 8F, 8G**, lower panels). These results support a role for CDK1 and Ki-67 in regulating microtubule and chromosome organization.

Moreover, we analyzed the distribution of nuclear lipids after RO-3306 treatment. Similar to the results of Ki-67 knockdown, the signal of nuclear lipids stained by DHPE-biotin was mostly smeared around the chromosomal area, including intrachromosomal regions, after RO-3306 treatment (**Figure 9A**). RO-3306 impaired the perichromosomal distribution of nuclear lipids at prometaphase (**Figure 9B**). The ratio of intra-to peri-chromosomal DHPE-biotin signals was also increased by the RO-3306 treatment (**Figure 9C**). These results further support that the key cell cycle regulator CDK1 regulates the distribution of nuclear lipids.

**Figure 9.**
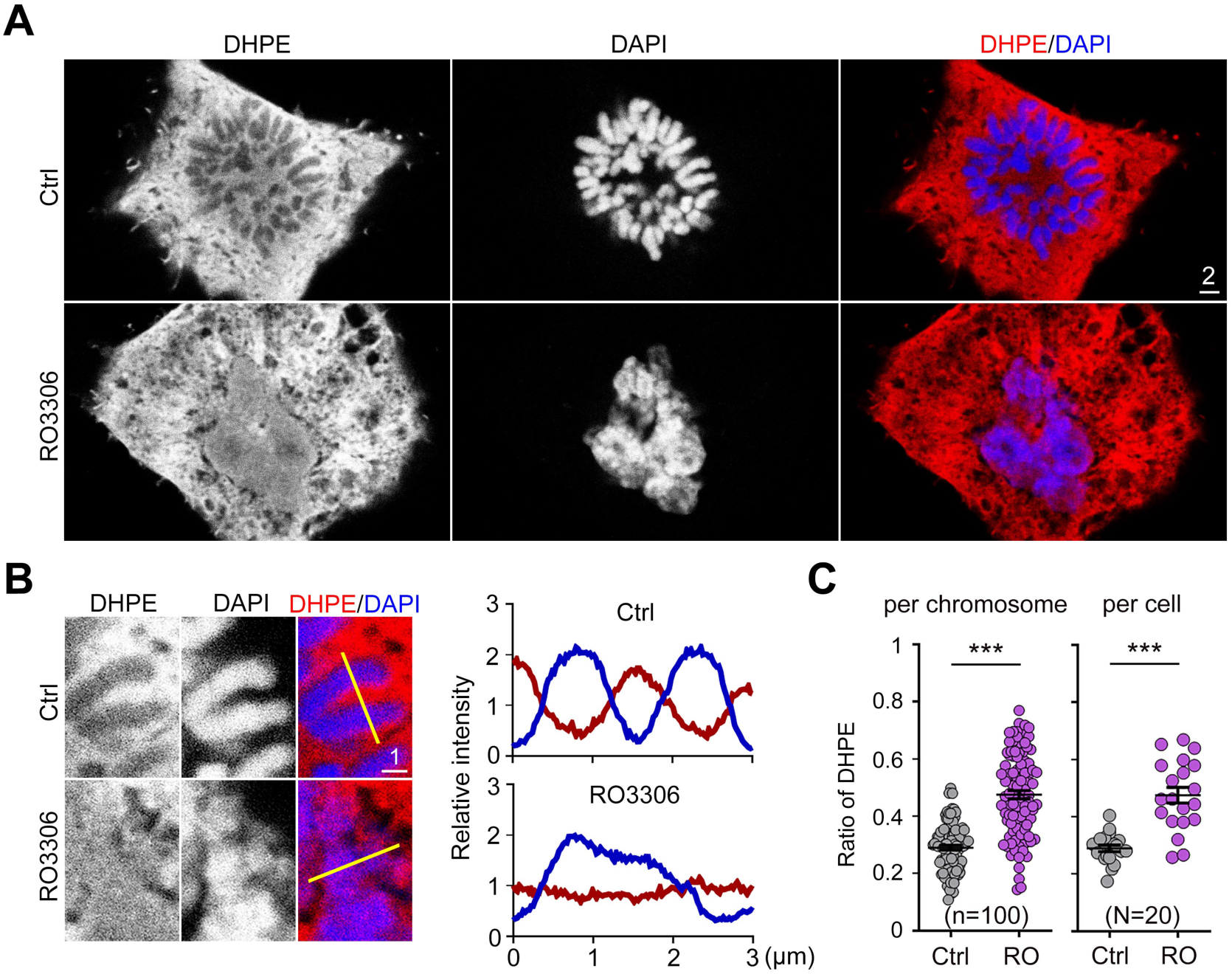
CDK1 inhibitor treatment alters nuclear lipid distribution. COS-7 cells were treated with RO-3306 and then stained with biotin-DHPE, Ki-67 and DAPI. An LSM700 microscope was used to acquire images of the cells in prometaphase. (A-C) Reduced Ki-67 phosphorylation alters lipid distribution patterns. (A) Representative confocal images of biotin-DHPE in Ctrl and RO-3306-treated cells during prometaphase. (B) Line scanning analysis of biotin-DHPE distribution across the chromosome. The yellow line indicates the region subjected to line scanning. (C) Ratio of biotin-DHPE signal in the intrachromosomal area to that at the periphery. RO-3306 treatment enhanced intrachromosomal signals. The sample sizes “N” or “n” indicate the number of examined cells and chromosomes, respectively. The data represent mean ± SEM and individual data points are also shown. All data were collected from two independent experiments. *** *P* < 0.001; unpaired two-tailed *t*-test. Scale bars: (A) 2 μm; (B) 1 μm; (D) 2 μm.

## Discussion

Here, we have applied a super-resolution fluorescence imaging system, i.e., TT-ExM (Wang et al., 2023, Hu et al., 2023), to reveal the dynamic distribution of nuclear lipids during the cell cycle, showcasing the involvement of nuclear lipids in how Ki-67 acts in chromosome individualization and segregation during mitosis. Our study provides three lines of evidence that Ki-67 regulates the distribution of nuclear lipids on the outer surface and/or interior of condensed chromosomes. First, although Ki-67 and nuclear lipids intermingle in the perichromosomal layer, our application of TT-ExM super-resolution imaging enabled us to dissect the layer organization at the chromosomal periphery, revealing that nuclear lipids or lipid-containing materials are enriched outside of Ki-67. Therefore, Ki-67 may act as a barrier to prevent nuclear lipids from moving into chromosomal interiors. Second, the enrichment of nuclear lipids at the chromosomal periphery is negatively correlated with total protein and phosphorylation levels of Ki-67 during mitosis. Third, Ki-67 knockdown results in nuclear lipids invading the interior of mitotic chromosomes, confirming that Ki-67 regulates the distribution of nuclear lipids during the cell cycle (**Figure 10**).

**Figure 10.**
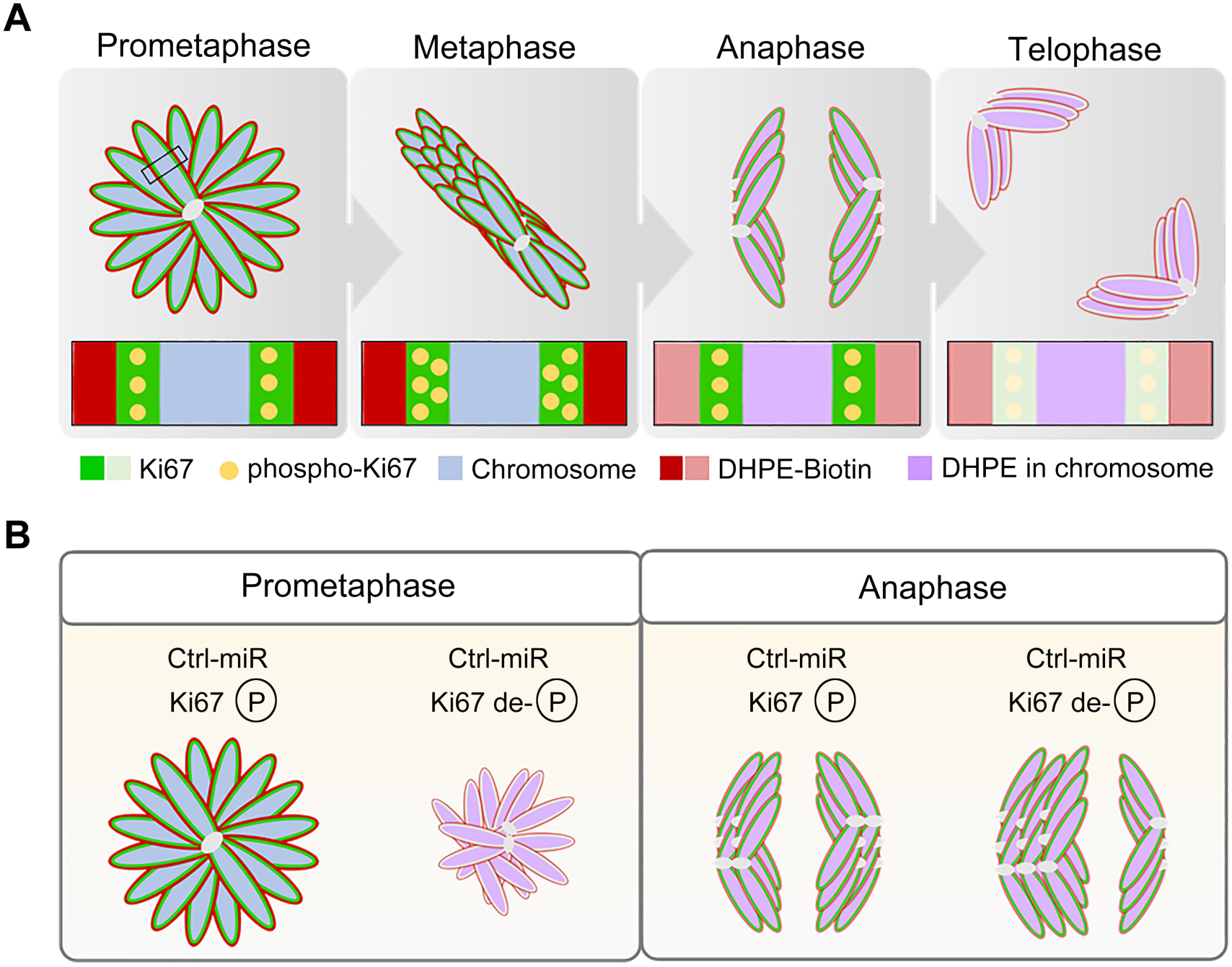
Dynamic alterations of nuclear lipids and Ki-67 phosphorylation and protein levels during mitosis. (A) Ki-67 regulates the distribution of nuclear lipids at the chromosomal periphery and forms the chromosomal envelope that regulates chromosome individualization during mitosis. (B) Ki-67 knockdown and Ki-67 dephosphorylation alter the lipid pattern at the chromosomal periphery and impair correct segregation of the sister chromatins.

How does Ki-67 regulate the distribution of nuclear lipids at the periphery of condensed chromosomes? It has been suggested previously that Ki-67 may serve as a surface-active agent (surfactant) to mediate the individualization of sister chromatid pairs (Cuylen et al., 2016) and to exclude the cytoplasm from fully condensed chromosomes (Cuylen-Haering et al., 2020). Although the respective mechanisms remain unclear, it is reasonable to speculate that nuclear lipids enriched at the chromosomal periphery likely cooperate with Ki-67 to facilitate the individualization of sister chromatid pairs during mitosis. The general physicochemical features of Ki-67 provide clues to the functional interplay between Ki-67 and nuclear lipids. Like many surfactants, Ki-67 has an amphiphilic structure. The long N-terminal domain of Ki-67 is excluded from chromatin, whereas its short C-terminal LR domain displays strong attraction to chromatin (Cuylen et al., 2016). Moreover, the chromosome separation function of Ki-67 is not attributable to a specific protein domain, but is correlated with the size and net charge of truncation mutants that lack secondary structure (Cuylen et al., 2016). Thus, the amphiphilic molecular structure of Ki-67 may disperse nuclear lipids or lipid-containing components via a liquid-liquid phase transition mechanism, resulting in their accumulation at the chromosomal periphery. Consistent with this supposition, our TT-ExM super-resolution imaging revealed that Ki-67 forms numerous tiny granules on the surface of mitotic chromosomes, rather than presenting a smooth and continuous thin layer. These tiny Ki-67-positive granules intermingle with lipid-containing nuclear components on the outer surface of mitotic chromosomes (**Figure 3A, 4D**). We suggest that Ki-67 proteins and these lipid-containing components collectively form a perichromosomal layer that mediates the individualization of sister chromatid pairs during mitosis.

In addition to forming a physical barrier via liquid-liquid phase separation (LLPS), CDK1-mediated Ki-67 phosphorylation may also regulate the distribution of nuclear lipids. We found that addition of the CDK1 inhibitor RO-3306 disrupted the organization of microtubules and chromosomes and induced nuclear lipid invasion into intrachromosomal regions at prometaphase. Heavy phosphorylation of Ki-67 may facilitate the formation of a negatively-charged barrier and strengthen the effect of Ki-67 in preventing nuclear lipid invasion into the intrachromosomal regions. However, since CDK1 phosphorylates many other proteins during mitosis (Petrone et al., 2016, Lo Surdo et al., 2023), other CDK1 substrates may also contribute to regulating the distribution of nuclear lipids. Nevertheless, our analyses using CDK1 inhibitors in the current study support that the cell cycle controls nuclear lipid distributions (**Figure 10**).

Since the lipid-containing nuclear components also display amphiphilic characteristics, it is also plausible that Ki-67 and the lipid-containing components may function redundantly, at least partially, thereby explaining several interesting observations. First, Ki-67 is not an essential protein but is a cell proliferation marker, with Ki-67 knockout cells still proliferating under cultured conditions (Cuylen et al., 2016). Second, to date, Ki-67 homologs have only been detected in vertebrates (Remnant et al., 2021). Proteins homologous to Ki-67 have not been identified or characterized in any other non-vertebrate eukaryotes. Our results indicate that nuclear lipids exert a novel role in isolating or individualizing condensed sister chromatid pairs during mitosis of cultured mammalian cells. It would be interesting to investigate further if nuclear lipids in non-vertebrate organisms also exhibit similar functions.

Our results are also consistent with previous reports (Prudovsky et al., 2012, Gould et al., 2017) showing that PS is mainly present in epichromatin or on the outer surface of condensed chromosomes during mitosis (**Figure 4**). Using biotin-DHPE, we show that nuclear lipids or lipid-containing components accumulate around the periphery of entire chromosomes during mitosis (**Figure 4**). Mammalian nuclei contain a variety of lipids, including PS, phosphatidylinositol, cholesterol, free fatty acids, diacylglycerol, and sphingolipids (Ledeen and Wu, 2004). Further investigations are needed to reveal if all these lipids or specific types of lipids are enriched around the periphery of condensed chromosomes.

Lastly, our findings herein may provide a new perspective on mitosis in general. The NE, which comprises two lipid bilayers, is a highly regulated membrane barrier that separates the nucleus from the cytoplasm in eukaryotes. Depending on the organism and cell type, the NE remains intact during closed mitosis or disassembles during open mitosis before chromosome segregation. Closed mitosis occurs in many (but not all) fungi and is considered the most ancient mechanism of eukaryotic cell division. In contrast, animals and plants undergo open mitosis, which appears to have evolved independently several times (Cavalier-Smith, 2010). A few fungi (e.g., the fission yeast *Shizosaccharomyces japonicus* and the corn smut fungus *Ustilago maydis*) undergo semi-open mitosis, in which the NE does not disassemble completely (Boettcher and Barral, 2013). As yet, it is not fully understood why open mitosis prevails in animals and plants, but not in fungi. The results from this study reveal that Ki-67 and nuclear lipids constitute a thin “chromosome envelope” around each sister chromatid pair, with the transition from “nuclear envelope” to “chromosome envelope” enabling continued isolation of genetic material from cytoplasmic material, thereby ensuring the individualization of sister chromatid pairs, the formation of star-like microtubule arrays, and proper chromosome segregation.

## Methods

### Antibodies and reagents

Rabbit polyclonal HSP90 was a kind gift from Dr. Chung Wang (Institute of Molecular Biology, Academia Sinica). Commercially available antibodies and chemicals used in this report are as follows: rat monoclonal Ki-67 (SolA15, #14-569882, 1:2000 for immunoblotting, 1:500 for TT-ExM, Invitrogen); rabbit polyclonal phospho-Ki-67 (301119, 1:500, Novocastra); mouse monoclonal phosphatidylserine (PS) (1H6, 05-719, 1:200 for TT-ExM, Sigma-Aldrich); rabbit polyclonal α-Tubulin (ab18251, 1:500, Abcam); chicken polyclonal GFP (ab13970, 1:1000 for TT-ExM, Abcam); mouse monoclonal GFP (JL-8, 632381, 1:2000 for immunoblotting, Clontech); Alexa Fluor-488-, 555- and 647-conjugated secondary antibodies (Invitrogen); Nocodazole (M1404, Sigma-Aldrich); DAPI (D1306,1:1000, Invitrogen); PicoGreen (P11495, 1:1000, ThermoFisher). Reagents used for DHPE TT-ExM are as follows: N-(Biotinoyl)-1,2-dihexadecanoyl-snglycero-3-phosphoethanolamine, triethylammonium salt (Biotin-DHPE) (60022, 0.1mg/ml, Biotium); VECTASTAIN ABC-HRP Kit, peroxidase (standard), (PK-4000, Vector Laboratory); Alexa fluor-555-conjugated tyramide (B40955, 1:100, Invitrogen); Amplification diluent (FP1050, PerkinElmer); 6-((acryloyl)amino)hexanoic acid (Acryloyl-X, SE, A-20770, ThermoFisher); Sodium acrylate (408220, Sigma-Aldrich); Acrylamide (A9099, Sigma-Aldrich); N, N’-methylenebisacrylamide, (1551624, ThermoFisher); Ammonium persulfate, (APS, A9164, Sigma-Aldrich); N,N,N,,N,-Tetramethylethylenediamine (TEMED, T7024, Sigma-Aldrich); 4-Hydroxy-TEMPO (176141, Sigma-Aldrich); Sodium chloride (3624-69, J.T.Baker); Trypsin-EDTA solution, 0.05% (25300-054, Gibco); RO-3306 (SML0569, Sigma-Aldrich); Methanol (9093-68; J.T. Backer); Acetone (32201, Honeywell).

### Plasmid construction

To generate the Ki67-miR knockdown constructs, an artificial miRNA was designed using BLOCK-iT™ RNAi Designer (https://rnaidesigner.thermofisher.com/rnaiexpress/). According to the ranking score, the top miRNA (nucleotide residues 3296-3319, AAGAGCTTGCCTGATACAGAA) was chosen and cloned into pcDNA6.2-GW/EmGFP vector. A control vector (pcDNA6.2–GW/EmGFP-miR, Invitrogen) expressing a scrambled miRNA (Hu et al., 2020) predicted to not target any gene in mammalian genomes was used as the negative control in our knockdown experiments.

### COS-7 culture and nocodazole treatment

The monkey COS-7 fibroblast-like cell line (ATCC, CRL-1651) was maintained in DMEM supplemented with 10% FBS, 1% penicillin/streptomycin and 2 mM glutamine. For mitotic cell cycle analysis, cells were treated with 50 nM nocodazole for 6 h to synchronize the cell cycle in prometaphase. Media were then washed out and the cells were allowed to recover for 15, 30, or 60 min to re-enter the mitotic cycle. Mitotic stages were identified based on chromosome patterning. For the Ki-67 dephosphorylation experiment, we treated nocodazole-synchronized cells with 2 μM of the CDK1 inhibitor, RO-3306, for 15 or 30 min after washing out nocodazole.

### Transfection and immunoblotting of COS-7 cells

COS-7 cells were transfected with the indicated plasmids using PolyJet (SL100688, SignaGen Laboratories) according to the manufacturer’s instructions. For Ki-67 knockdown analysis, the transfection period was extended to 2 days to enhance knockdown efficiency. For immunoblotting analysis, cells were washed with PBS twice and lysed directly with SDS sample buffer. Total cell lysates were separated by means of SDS-PAGE and analyzed by immunoblotting.

### Immunofluorescence analysis

Regular immunostaining was performed as described previously (Hu et al., 2020). Briefly, cells were washed with phosphate-buffered saline (PBS) and fixed with 4% paraformaldehyde (PFA) and 4% sucrose in PBS for 10 min, followed by permeabilization with 0.2% Triton X-100 in PBS for 15 min at room temperature. After blocking with 10% bovine serum albumin (BSA), the cells were incubated with primary antibodies diluted in PBS containing 3% BSA and 0.1% horse serum at 4 °C overnight. After washing with PBS several times, the cells were incubated with secondary antibodies conjugated with Alexa Fluor-488, -555 or -647 (Invitrogen) and DNA dye for 2 h at room temperature.

### Lipid staining

Lipid labeling using biotin-DHPE (Wang et al., 2023) was performed before regular immunostaining. After fixation with 4% paraformaldehyde (PFA) for 10 min, the cells were washed with PBS and then treated with 0.1 mg/ml biotin-DHPE in 50% ethanol overnight at room temperature. After washing with PBS several times, cells were further incubated with premixed reagents A and B in the Vectastin ABC-HRP kit (Vector Laboratory) in TBS (1:50 dilution) for at least 30 min. After washing with PBS, Alexa fluor-555-conjugated tyramide was diluted (1:50) in amplification diluent and applied to the cells for 15 min. Following extensive washing, the biotin-DHPE-stained cells were then deemed ready for regular immunostaining. To confirm that the labeling of biotin-DHPE was lipid-dependent, 100% methanol or 100% acetone were added separately into PFA-fixed cells for 10 min at 20 °C. Comparing the results from PFA fixation alone with PFA followed by methanol or acetone treatment, we demonstrated lipid-specific staining by biotin-DHPE.

### TT-ExM

Immunostained COS-7 cells, prepared as described above, were subjected to TT-ExM (Wang et al., 2023, Hu et al., 2023). After staining, the cells were washed with PBS three times and then incubated with 0.1 mg/ml Acryloyl-X, SE in PBS overnight at 4 °C. After washing three times with PBS, the coverslip with stained cells was placed on acrylamide solution (with the cell-attached side facing down) at 37 °C for 2 h for gelation. After gelation was completed, the hydrogel was immersed in trypsin digestion solution (0.05 % volume dilution: 2 μl trypsin-EDTA solution in 4 ml low salt digestion buffer), and then incubated at room temperature for at least 20 h with gently shaking. The digested hydrogel was transferred to a 10-cm dish filled with excess Milli-Q water for free expansion for 2 h. The expanded samples were then imaged at room temperature using an LSM980 laser scanning confocal microscope equipped with Airyscan 2 multiplex mode SR-4Y. An LD LCI Plan-Apochromate 40X/1.2 Imm Corr DIC M27 objective was used for Z-series image acquisition.

### Analyses of DHPE invasion, chromosomal layer patterning and chromosome area

To quantify the immunoreactivities of DHPE signals in the intrachromosomal area, line scanning was performed using the ImageJ/Fiji software. A 3 µm-wide line across the chromosome or perichromosomal layer was drawn to quantify DHPE signals. The intersection of biotin-DHPE signal and the chromosome boundary was set as a ratio of 1. The lowest ratio of biotin-DHPE was identified as the lipid level in the intrachromosomal area. Ten individual chromosomes per cell were quantified and averaged. For chromosomal layer analysis, post-expanded and LSM980-acquired images were subjected to line scanning analysis across the perichromosomal layer. Line histograms were used to exhibit the distribution pattern of pericromosomal components. Five individual chromosomes per cell were quantified and averaged. For chromosome area analysis, the area occupied by chromosomes was quantified based on DAPI signals using ImageJ/Fiji. For chromosome segregation in anaphase, the total chromosome area and difference in area between the two sister chromosomes were analyzed.

### Analysis of microtubule organization

To analyze microtubule organization, cells were stained with α-tubulin to identify microtubule filaments and then assessed by means of Sholl analysis (ImageJ/Fiji) (Chen et al., 2011, Hu et al., 2020) to examine the extension pattern of tubule filaments and to count the numbers of intersections. The total numbers of intersections and microtubule filament tips were counted to represent microtubule organization. For the microtubule organization analysis, microtubules were categorized into four types based on distribution patterns, i.e., organized, disorganized, two-star, or extended. The percentage of each type was calculated to reveal the effect of RO-3306 on microtubules during mitosis.

### Quantification and statistical analysis

To avoid potential personal bias, all data were collected and analyzed blindly by having other lab members relabel the samples. For each experiment, 10 cells were randomly collected from at least two independent experiments for quantification. To analyze DHPE invasion into the chromosomal area, line scanning analysis was performed at each mitotic stage using the ImageJ/FIJI software. To analyze microtubule organization, Sholl analysis was performed in the ImageJ/FIJI software to calculate numbers of microtubule filaments and their distribution patterns. Two-group comparisons were analyzed with two-tailed unpaired Student’s *t*-test. For more than two group comparisons, a one-way analysis of variance (ANOVA) with Bonferroni’s multiple comparison post-hoc test was performed.

## Declarations

### Ethics approval and consent to participate

Not applicable.

### Consent for publication

All authors have read and approved of its submission to this journal.

### Availability of data and materials

All relevant data are within the paper and its Supporting information files. The file of **Supplemental Table S1** contains the numerical data for all figures. **Supplemental Table S2** contains all statistical results. **Supplemental Figure S1** contains uncropped blots of **Figure 5B**.

## Authors’ contributions

Conceptualization: YPH, TFW; Funding acquisition: YPH, TFW; Investigation: HTH, UTTW, BCC, YPH; Methodology: HTH, UTTW, BCC, YPH, TFW; Project administration: YPH, TFW; Supervision: YPH, TFW; Visualization: HTH, YPH; Writing – original draft: HTH, YPH, TFW; Writing – review and editing: HTH, UTTW, BCC, YPH, TFW. All of the authors read and approved the manuscript.

## Acknowledgements

We thank the Imaging Core Facility of the Institute of Molecular Biology, Academia Sinica, for technical support and John O′Brien for English editing.

## Authors’ information

Hsiao-Tang Hu

Ueh-Ting Tim Wang

Bi-Chang Chen

Yi-Ping Hsueh

Ting-Fang Wang

## Supplemental Information for

**Supplemental Figure S1.**
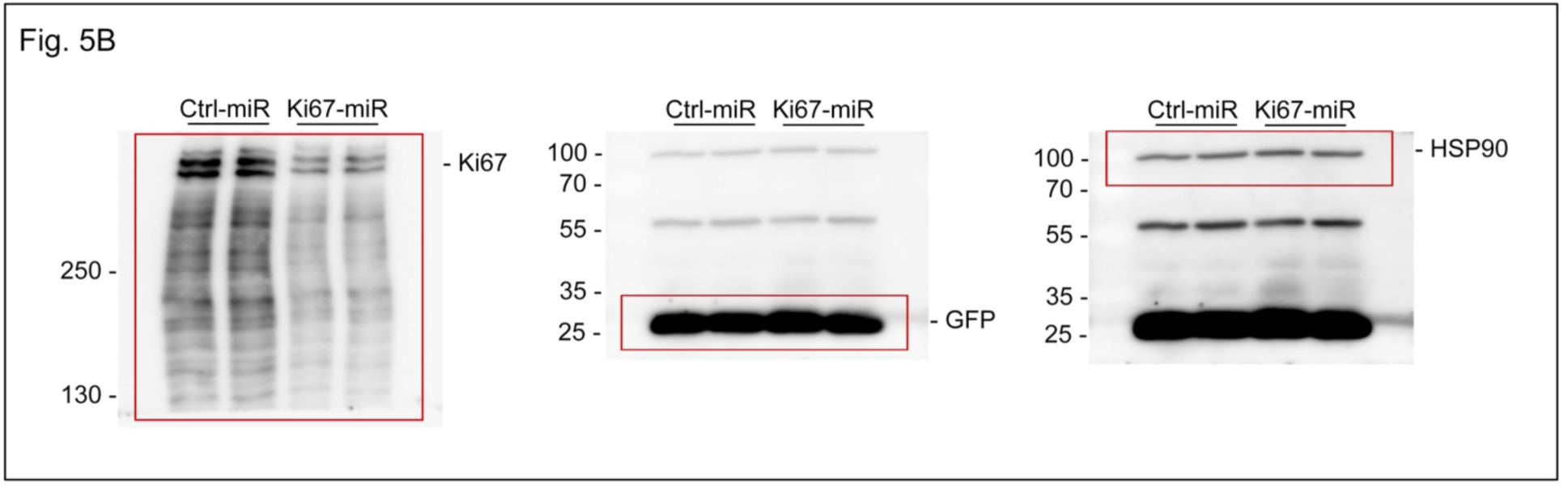
Uncropped immunoblots of. **Figure 5B**.

**Supplemental Table S1. The numerical data for all figures.**

**Supplemental Table S2. All statistical results.**

