## Supplementary figures and images for "Ki-67 and CDK1 Control the Dynamic Association of Nuclear Lipids with Mitotic Chromosomes"

### Supplemental Figure S1

**Supplemental_Fig_ S1. Uncropped immunoblots of Figure 5B.**

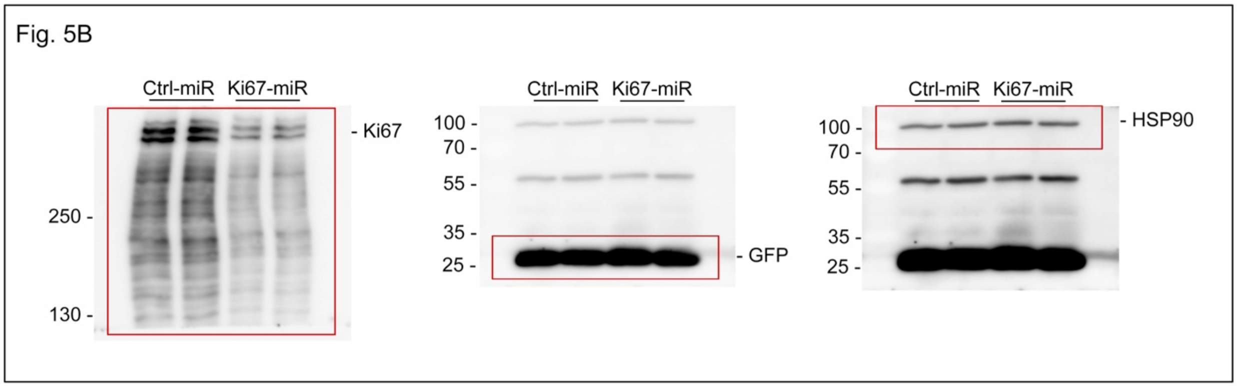
